# The River Runs Through It: the Athabasca River Delivers Mercury to Aquatic Birds Breeding Far Downstream

**DOI:** 10.1101/440115

**Authors:** Craig E. Hebert

## Abstract

This study examined factors contributing to temporal variability (2009-2017) in total mercury (THg) concentrations in aquatic bird eggs collected in the Peace-Athabasca Delta and Lake Athabasca in northern Alberta. Factors examined included annual changes in oil sands production, bird diets, forest fires, and flow of the Athabasca River. Surface mining activities associated with Alberta’s Athabasca oil sands are centered north of Fort McMurray, Alberta, adjacent to the northward-flowing Athabasca River. Previous studies have found that oil sands industrial operations release mercury into the local (within ~50 km) environment. However, temporal trends in egg THg levels did not track trends in synthetic oil production from the oil sands. Intraspecific fluctuations in bird diet also could not explain annual variability in egg THg levels. Annual extent of forest fires in Alberta was only related to egg THg concentrations in California Gulls from Lake Athabasca; annual levels in other species showed no relationship with fire extent. The inclusion of more terrestrial foods in gull diets may have contributed to this difference. For the majority of species, annual fluctuations in maximal flow of the Athabasca River were important in influencing annual egg THg levels. Eggs collected following years of high flow had higher THg concentrations with distinct stable Hg isotope compositions. Riverine processes associated with suspended sediment were likely critical in regulating Hg availability to nesting birds. This study highlights the importance of the Athabasca River as a conduit for Hg transport to ecologically-sensitive downstream ecosystems such as the Peace-Athabasca Delta and Wood Buffalo National Park (a UNESCO World Heritage Site). Human activities that increase atmospheric Hg deposition to the Athabasca River watershed, or that enhance Hg releases to the river through erosion of Hg-bearing soils, will likely increase the availability of Hg to organisms inhabiting downstream areas.

## Introduction

The Athabasca River is a major river flowing northeast 1200 km from the Rocky Mountains to Lake Athabasca. Along the way, it passes through Alberta’s oil sands, a region of large-scale, open-pit mines used to extract bitumen for synthetic oil production. The river discharges into the Peace-Athabasca Delta (PAD) and western Lake Athabasca approximately 200 km downstream of Fort McMurray (see [1]). Since 1967, when oil sands operations commenced, industrial exploitation of the bitumen-rich oil sands has expanded greatly [2] (Fig. S1). Previous research has indicated that industrial development associated with the oil sands is a source of mercury (Hg) to the environment [3, 4]. For example, Kirk et al. [4] found that deposition of both total mercury (THg) and methyl mercury (MeHg) in snow resembled a bullseye pattern with higher Hg levels in snow collected closer to oil sands developments. However, snow Hg levels declined rapidly with distance from such developments with most Hg being deposited within approximately 50 km. These studies [3, 4] highlighted the possibility that spring snowmelt could result in the release of chemicals, such as Hg, into the aquatic environment. Similarities in relative concentrations of metals in snow and river water provided evidence of metals emitted to the air finding their way into the Athabasca River and tributaries [3]. Kelly et al. [3] also found that Hg concentrations in water were greatest downstream of areas disturbed by oil sands development. Passage of water through the surface-mineable oil sands region, in itself, was not responsible for elevated water Hg levels. Degree of land disturbance caused by oil sands operations was important in enhancing water Hg concentrations. The significance of these findings in terms of effects on Hg concentrations in biota inhabiting areas farther downstream, e.g. the PAD, has not been documented and deserves further study.

The PAD is a wetland of international significance [5] and it is a defining feature of Wood Buffalo National Park (WBNP), a UNESCO World Heritage Site. It provides habitat for millions of birds and WBNP is the only breeding area for the endangered Whooping Crane (*Grus americana*) [6]. The surrounding region also provides important wildlife habitat. For example, Egg Island in western Lake Athabasca is a provincial ecological reserve harboring a variety of colonially-nesting aquatic birds including the largest breeding colony of Caspian Terns (*Hydroprogne caspia*) in Alberta [7]. The PAD and Lake Athabasca are also important to Indigenous land users who rely on traditional wild foods [8]. Hence, environmental changes brought about by human activities have the potential to significantly affect wildlife and human inhabitants of this region.

Colonial aquatic birds, e.g. gulls and terns, are commonly used to monitor the state of the environment [9]. These birds are top predators resulting in their accumulation of high levels of biomagnifying contaminants. Eggs are a useful sampling matrix for contaminant studies as their collection has little impact on bird populations and eggs of these species are formed from exogenous, locally-obtained resources [10]. Since 2009, gull and tern eggs have been collected annually from sites in WBNP (Mamawi Lake) and western Lake Athabasca (Egg Island). The goal of this study was to examine temporal trends (2009-2017) in egg THg levels and determine whether they were influenced by a variety of factors including: oil sands production, bird diet, forest fires, and flow of the Athabasca River. We briefly discuss each of these factors below.

From 2009-2016, oil production from surface mineable sources in the Athabasca oil sands increased by 47% (761,000 bpd (barrels per day) to 1,117,000 bpd) and accounted for approximately 46% of total oil sands production in 2016 (in-situ production accounted for the remainder) [2]. As oil sands operations are a known source of Hg [3, 4], it is plausible that increases in oil/bitumen production might be accompanied by increases in Hg releases to the environment that could result in increased Hg levels in wildlife receptors such as bird eggs. Because MeHg biomagnifies, it is possible that annual changes in bird diets, particularly with respect to bird trophic position, could be important in regulating annual egg THg concentrations. Inter-specific differences in food sources could also be important in regulating Hg exposure. Forest fires are known to release Hg into the environment [11, 12] and Alberta experienced very large forest fires in some years during the study period [13]. It is possible that such fires could increase Hg levels in the food webs used by birds with resultant increases in egg THg concentrations. Previous research on aquatic birds has highlighted the possibility that the Athabasca River may be a source of Hg to wildlife inhabiting downstream environments [14]. In a large-scale spatial assessment of Hg levels in western Canadian gull eggs, Dolgova et al. [15] found that levels were greatest in eggs collected from breeding sites in receiving waters of the Athabasca River. Hence, factors associated with the regulation of Hg transport via the Athabasca River may be important in regulating Hg levels in downstream wildlife. One such critical factor is annual maximal river flow. Long and Pavelsky [16] documented the influence of river flow on sediment transport down the Athabasca River into the PAD and Lake Athabasca. In the river, there was a linear relationship between flow and suspended sediment concentrations (SSC) with SSC notably increasing around flows of 1500 m^3^/s. In addition, a non-linear relationship between river flow and downstream SSC in western Lake Athabasca was observed after maximal flow exceeded a threshold of 1700 m^3^/s. Riverine transport of sediment-bound Hg is an important mechanism for the transfer of that element to downstream environments [17]. SSC may also influence Hg dynamics in the environment through effects on the photochemical reduction of MeHg. These effects, and possible Hg sources, can be evaluated using Hg stable isotopes.

Mass-dependent (MDF) and mass-independent (MIF) fractionation of stable Hg isotopes (^196^Hg, ^198^Hg, ^199^Hg, ^200^Hg, ^201^Hg, ^202^Hg, ^204^Hg) can provide insights into processes regulating Hg dynamics in the environment. MDF, e.g. δ^202^Hg, is influenced by a variety of factors including physical processes as well as by biological reactions [18] and can be useful in identifying the flow of mercury from terrestrial and aquatic systems to consumers [19]. MIF, on the other hand, has largely been documented for the odd mass Hg isotopes (^199^Hg, ^201^Hg) and is thought to primarily be the result of a magnetic isotope effect, i.e. dissimilarities in magnetic spin of even and odd mass isotopes result in their reacting at different rates. Existing evidence suggests that MIF is not influenced by food web/trophic interactions [20] but occurs during photochemical MeHg degradation and photoreduction of Hg^2+^ [21]. Therefore, factors that affect Hg exposure to light, e.g. light penetration through the water column, are expected to influence MIF of Hg isotopes [22]. Photodegradation of MeHg to Hg^0^ may be an important MeHg removal mechanism, particularly in systems characterized by clear water with a high degree of light penetration, and this can be assessed using Hg isotopes [23]. However, only a portion of the MeHg in such systems will undergo photochemical demethylation. The remaining MeHg will exhibit MIF of ^199^Hg and ^201^Hg and that MeHg may be incorporated into food webs with Hg isotope values in consumers (e.g. birds) reflecting MIF of Hg isotopes [23].

The slope of the MIF of ^201^Hg versus ^199^Hg is useful in differentiating the Hg species (MeHg or Hg^2+^) undergoing photoreduction (MeHg slope ~ 1.3, Hg^2+^ slope ~1.0) [21, 23, 24]. A slope near 1.3 indicates that photoreduction of MeHg is the main mechanism underlying MIF values. If photochemical reduction of MeHg is the main process underlying MIF of Hg isotopes then Δ^199^Hg and/or Δ^201^Hg values in consumer tissues can be used to estimate the amount of MeHg that has been photodegraded [21, 25].

In this study, annual THg levels in eggs of aquatic birds nesting downstream of the Athabasca River are assessed. As described above, various factors, i.e. oil sands production, bird diet, forest fires, and riverine processes, are investigated to determine the degree to which they explain patterns in annual egg THg concentrations. As part of this analysis, the degree to which maximal annual flow of the Athabasca River and concomitant changes in SSC are important in regulating Hg availability to birds is investigated. Mercury isotopes are used to gain insights into the processes regulating Hg dynamics and availability in this ecosystem.

## Methods

### Field Methods

Aquatic bird, i.e. gull and tern, egg collections were made in June, 2009-2017 (no collections were made in 2010) at two sites located in receiving waters of the Athabasca River. Mamawi Lake (58.60N, -111.47W) is located in the Peace-Athabasca Delta within the boundaries of Wood Buffalo National Park. Egg Island (59.98N, -110.44W) is located in the western end of Lake Athabasca approximately 40 km northeast of the mouth of the Athabasca River (see Dolgova et al. [15] for map showing collection locations). In most years, 10 eggs were collected per species per site. At Mamawi Lake, Ring-billed Gull (*Larus delawarensis*) and Common Tern (*Sterna hirundeo*) eggs were collected. Low water levels in 2011 prevented access to that location so no eggs were collected that year. Also, in 2014 it was impossible to collect Common Tern eggs at that site. At Egg Island, 10 eggs were collected annually from three species: California Gull (*Larus californicus*), Caspian Tern, and Common Tern. The first recorded nesting of Common Terns on Egg Island was in 2011 so eggs of that species were not collected from that site in 2009. Eggs were transported to the National Wildlife Research Centre, Ottawa, ON, Canada in padded cases.

In 2013, prey fish, i.e. cyprinids, were collected from Mamawi Lake and two sites in western Lake Athabasca to provide isotope data for bird prey. Whole-body fish were analyzed of two surface-schooling species: Spottail Shiner (*Notropis hudsonius*) and Emerald Shiner (*N. atherinoides*). These species are accessible to surface-feeding birds. Details regarding prey fish collections are available in Dolgova et al. [15].

Epiphytic lichen was collected from the branches of coniferous tree species in 2013 (May or July) from 14 locations north, south, east and west of Fort McMurray. All sites were within 100 km of that city. Samples (n=26, two samples per site except at two sites where only one sample was analyzed) were cleaned by removing foreign debris, freeze dried, and pulverized into a fine powder before analysis. The utility of tree lichens as passive collectors of atmospheric pollutants, including mercury, has been demonstrated [26, 27]. Mercury isotope measurements in lichens were used to establish baseline Hg isotopic “signatures” associated with atmospherically deposited mercury in the Fort McMurray region. This would include mercury from all sources to that region including the oil sands.

### Laboratory Methods

Egg contents (i.e. yolk, albumen, embryonic tissue) and whole-body prey fish were homogenized/pulverized using liquid nitrogen and a cryogenic ball-mill. Homogenates were aliquoted into acid-washed glass and polypropylene containers and stored at −40° C prior to analysis.

Details regarding egg THg analysis are identical to those reported in Hebert and Popp [28]. In eggs, approximately 97% of THg is in the organic MeHg form [29] so measuring THg is a cost-effective way to assess MeHg levels in eggs. THg concentrations measured in all samples were within concentration ranges of certified reference materials (OT1566b, TORT-3, DOLT-4, IAEA-407, BCR-463). Limit of detection for THg was 0.006 μg/g (dry weight).

Stable isotopes of nitrogen (^15^N and ^14^N expressed as δ^15^N) and carbon (^13^C and ^12^C expressed as δ^13^C) were measured in prey fish (collected in 2013) and in the contents of individual eggs (all years) to assess bird diets across years. Details regarding stable isotope analysis are identical to those reported in Hebert and Popp [28]. Dietary change affecting the trophic position of laying females is reflected in egg δ^15^N values (‰) as nitrogen isotopes undergo MDF. MDF leads to enrichment of ^15^N in higher trophic level organisms with increasing δ^15^N values with increasing trophic position. Because Hg biomagnifies, temporal changes in trophic position would be expected to affect female exposure to, and uptake of, MeHg with concomitant effects on egg THg levels. Carbon isotopes are useful in evaluating carbon sources utilized by consumers [30]. Because lipid content of samples can affect δ^13^C values, a mathematical approach was used to adjust δ^13^C values in eggs and prey fish based upon C:N ratios in individual samples (C:N ratios for all samples were > 4.0). For eggs, the equation used was from Elliott et al. [31], *δ*^13^C_lipid-corrected_ = *δ*^13^C_non-corrected_ − 4.46 + 7.32 * Log (C:N ratio). For fish, *δ*^13^C_lipid-corrected_ = *δ*^13^C_non-corrected_ − 3:32 + 0:99 * C:N [32].

Mercury isotope analysis was conducted at Trent University’s Water Quality Centre. Details regarding mercury isotope analysis have been reported previously [33]. Mass-independent fractionation (MIF) of Hg isotopes is unrelated to isotope mass and is reported in capital delta (Δ) notation. MIF describes the difference between measured δ^xxx^Hg values and scaled δ^202^Hg values (Δ^xxx^Hg = δ ^xxx^Hg − (δ^202^Hg × β)). β, the scaling factor, is determined by theoretical laws of mass dependent fractionation (MDF) and is an isotope-specific constant. Here, the focus is on the interpretation of MDF (δ^202^Hg) and MIF results (Δ^199^Hg and Δ^201^Hg). MDF data can provide insights into pathways of mercury exposure, e.g. terrestrial versus aquatic [19], while MIF can be useful in assessing the processes influencing mercury isotope values in eggs. Hg isotope analyses were conducted on eggs from all four study species from both study sites allowing an assessment of whether MIF of Hg isotopes was influenced by bird trophic position (as inferred from egg δ^15^N values; see below). Samples were analyzed from a variety of years (2009-2017) for the different species.

### Analytical and Statistical Methods

Annual estimates of synthetic oil/bitumen production from surface-mined oil sands deposits were obtained from CAPP [2].

Annual estimates of total forest fire extent (ha) for the province of Alberta were obtained from the National Forestry Database [13]. During the period of study, 99% of total annual area burned occurred during May-August with June-August usually being the focal months. Hence, these fires would have occurred after the egg-laying period in that same calendar year. Caldwell et al. [34] reported sediment MeHg levels peaked within three months after a wildfire, further delays would be expected in terms of wildfire-generated Hg being incorporated into food webs. Therefore, fire extent in the year preceding egg collection was used to assess the impact of fire on egg THg levels. The one-year time lag between forest fire extent and egg THg levels allowed time for Hg released from fires to be incorporated into, and passed through, the food webs utilized by birds.

Annual monthly estimates of Athabasca River flow were obtained from a hydrometric station located five km north of Fort McMurray (station 07DA001) (https://wateroffice.ec.gc.ca/mainmenu/historical_data_index_e.html). During the period of study, maximal monthly river flows were most often observed in June (seven of 10 years from 2008-2016), therefore, June flow was used to estimate maximal annual river flow. Based upon Long and Pavelsky [16], years were categorized as being low (June mean monthly flow <1600 m^3^/s) or high (June mean monthly flow ≥1600 m^3^/s) flow years. River flow in the year preceding egg collections was used to categorize egg collections as being influenced by low or high flow years. For example, mean river flow in June 2011 was >1600 m^3^/s, hence egg collections made in June 2012 were categorized as being influenced by a high flow year. The one-year time lag between river flow and egg THg levels allowed for the influence of river processes on Hg availability/dynamics to be incorporated into downstream food webs. This period of time is consistent with work examining the rapidity with which Hg is incorporated into foodwebs. For example, mercury levels in small fish responded within 1-2 years to changes in levels of Hg in the environment [35]. Such small fish species are an important component of the diet of gulls and terns and because eggs of these species are formed from locally-obtained exogenous resources, e.g. prey fish, we would expect egg Hg levels to respond within a similar time-frame to prey fish. For statistical comparisons, years were combined into either low or high flow categories and these categories were compared.

True color Landsat images [36] of the PAD and western Lake Athabasca were obtained from years of low (2002, 2010, 2015) and high (2011, 2014, 2017) Athabasca River flow. These images were used to qualitatively visualize annual differences in the extent of sediment plumes entering western Lake Athabasca.

Data regarding suspended sediment concentrations (SSC) and secchi depth (a measure of water clarity) were obtained from Long and Pavelski [37]. Data were available from sampling sites situated along a southeast-northwest transect crossing the western basin of the lake (see [16]). Distance of each sampling site from the mouth of the Athabasca River was estimated based upon georeferenced data. Data from low (2010) and high (2011) flow years were used to assess inter-year differences in the influence of the river on SSC in western Lake Athabasca and resultant impacts on water transparency.

Inter-year differences in variables were assessed using ANOVA or Kruskal-Wallis tests followed by selected post-hoc tests (Tukey’s HSD or Dunn’s test). Relationships between variables were evaluated using Pearson correlation coefficients (*r*). Student’s two-tailed t-tests were used to compare egg THg levels and Hg isotope values between years of low and high Athabasca River flow. All statistical analyses were conducted using Statistica Ver 12 [38] with α = 0.05. Assumptions underlying the use of parametric statistics were tested using Q-Q plots, the Shapiro-Wilk W test (normality), and Levene’s test (homogeneity of variances).

## Results

### Temporal trends in egg THg concentrations (2009-2017)

Egg THg concentrations showed inter-year differences for most species/site combinations (Egg Island: Caspian Terns ANOVA *F*(7,72) = 8.83, *p* < 0.001; Common Terns ANOVA *F*(6,63) = 8.96, *p* < 0.001; Mamawi Lake: Ring-billed Gulls Welch’s ANOVA *F*(6,30) = 3.61, p = 0.01; Common Terns Welch’s ANOVA *F*(5,20) = 3.92, *p* = 0.01) except for Egg Island California Gulls (Welch’s ANOVA *F*(7,31) = 0.89, *p* = 0.53). However, egg THg concentrations were not correlated with year of collection for Egg Island Caspian Terns (n = 80, *r* = −0.01, *p* = 0.97), Egg Island Common Terns (n = 70, *r* = 0.08, *p* = 0.53), or Mamawi Lake Ring-billed Gulls (n = 83, *r* = −0.13, *p* = 0.24) (Figure 1A, B). THg levels in eggs of Egg Island California Gulls increased through time (n = 81, *r* = 0.24, *p* = 0.03) while THg levels in Common Tern eggs from Mamawi Lake decreased (n = 55, *r* = −0.40, *p* = 0.002) (Fig.1*A*, *B*). In both these cases, temporal trends were influenced by data from one year. For the Egg Island California Gull eggs, high THg levels in 2017 were responsible for the temporal increase. When 2017 data were removed from the analysis, there was no significant relationship with time (n = 71, *r* = 0.08, *p* = 0.52).

**Fig 1.**
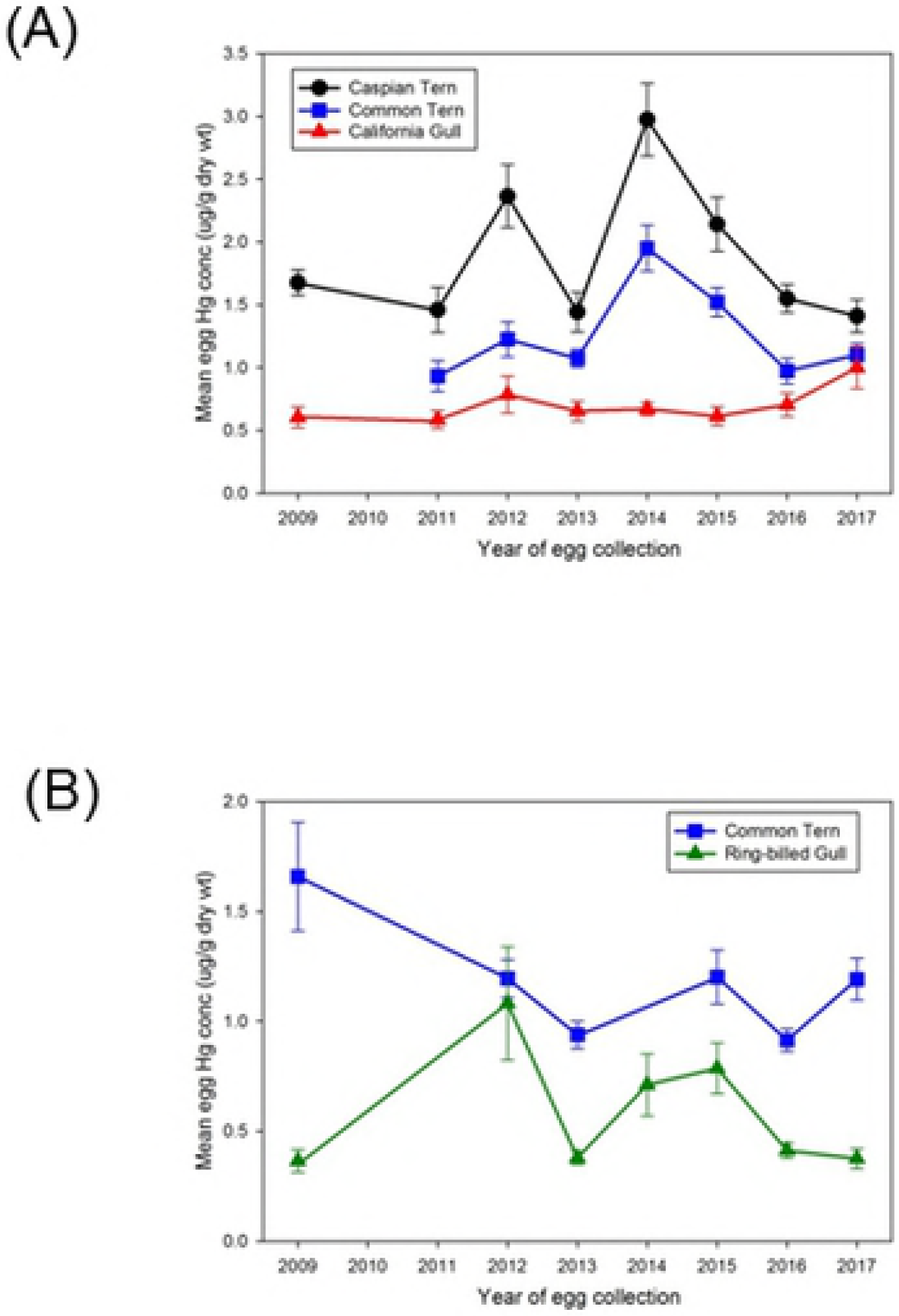
Annual mean (± SE) THg levels (ug/g, dry weight) in gull and tern eggs from sites in receiving waters of the Athabasca River. (a) Egg Island, western Lake Athabasca: Caspian Terns, Common Terns, California Gulls (b) Mamawi Lake: Common Terns, Ring-billed Gulls.

Similarly, the declining temporal trend in THg in Common Tern eggs from Mamawi Lake was driven by the 2009 data. When data from that year were omitted from the analysis, no temporal trends were evident (n = 45, *r* = −0.06, *p* = 0.71).

### Influence of oil sands production, forest fires, and bird diet on egg THg

Temporal trends in egg THg concentrations were not related to annual production of synthetic oil/bitumen from surface-mined oil sands deposits for any of the species/site combinations (2009-2016 annual data. Egg Island CAGU n = 8, *r* = .56, *p* = .15; CATE n = 8, *r* = −.03, *p* = .94; COTE n = 7, *r* = .07, *p* = .89; Mamawi Lake RBGU n = 7, *r* = −.13, *p* = .78; COTE n = 6, *r* = −.67, *p* = .14). The production of synthetic oil/bitumen from surface mining showed a significant increase during the period of study (*r* = 0.996, *p* < 0.001) (Fig. S1) which was not reflected in the annual fluctuations observed in egg THg levels (Fig.1*A*, *B*).

Annual extent of forest fires preceding the year of egg collection showed no relationship with THg levels in Mamawi Lake eggs (Ring-billed Gulls, *r* = 0.23, *p* = 0.59; Common Terns *r* = −0.46, *p* = 0.36) nor in eggs of Caspian Terns (*r* = −0.18, *p* = 0.64) or Common Terns (*r* = −0.54, *p* = 0.16) at Egg Island. However, mean annual egg THg levels were correlated with forest fire extent in eggs of California Gulls (*r* = 0.73, *p* = 0.03) at that location (Fig. 2).

**Fig 2.**
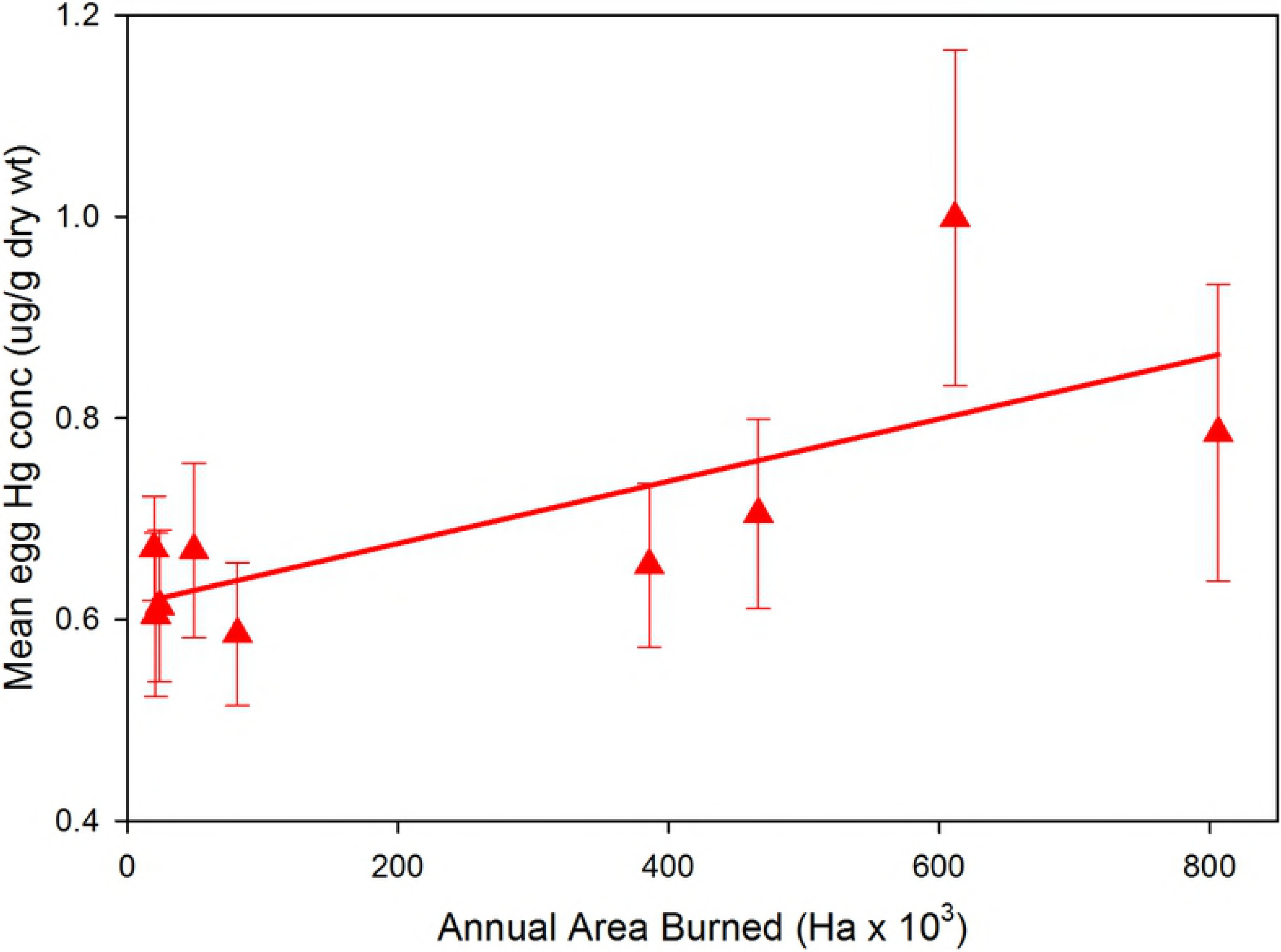
Annual extent (ha) of forest fires in Alberta versus annual mean (± SE) THg levels (ug/g, dry weight) in California Gull eggs from Egg Island, Lake Athabasca. Annual egg THg levels were compared to forest fire extent in the year preceding egg collections.

Egg stable nitrogen isotope values (δ^15^N) were different among all species comparisons (Kruskal-Wallis *H*(3,369) = 171.85, *p* < 0.001, Dunn’s test, Caspian Tern > Common Tern > California Gull > Ring-billed Gull). δ^15^N values within most species and sites showed no differences across years (Egg Island: California Gulls ANOVA *F*(7,73) = 2.12, *p* = 0.051; Caspian Terns ANOVA *F*(7,72) = 1.09, *p* = 0.38; Common Terns Kruskal-Wallis *H*(6,70) = 9.37, *p* = 0.15; Mamawi Lake: Ring-billed Gulls Kruskal-Wallis *H*(6,83) = 12.86, *p* = 0.05, Dunn’s post-hoc test no differences). Only Mamawi Lake Common Terns showed significant inter-year differences in egg δ^15^N values (Kruskal-Wallis *H*(5,55) = 20.76, *p* = 0.01, Dunn’s test) (Table S1). For three of the five species/site combinations, inter-year differences in δ^13^C values were observed but these were minimal with most years having similar values (Table S2).

Interspecific comparisons of stable Hg and carbon isotope values revealed significant differences among species (Fig.3). Egg δ^202^Hg values reflecting MDF of Hg isotopes were greater in California Gulls than the other bird species (ANOVA *F*(3,185) = 100.0, *p* < 0.001, followed by Tukey’s HSD test). Egg δ^13^C values were less negative in California Gulls (mean = −23.7‰) than the other three species (mean values; Common Tern = −26.3‰, Caspian Tern = −25.8‰, Ring-billed Gull = −25.7‰). δ^13^C values in Ring-billed Gulls were also less negative than Common Terns (Welch’s ANOVA *F*(3,185) = 91.1, *p* < 0.001, followed by Tukey’s HSD test). δ^13^C values in California Gulls showed the greatest deviation from those in prey fish (mean ± 1 SD values, Emerald Shiner = −28.33 ± 2.10 (n=46), Spottail Shiner = −28.00 ± 3.41 (n=22)).

**Fig 3.**
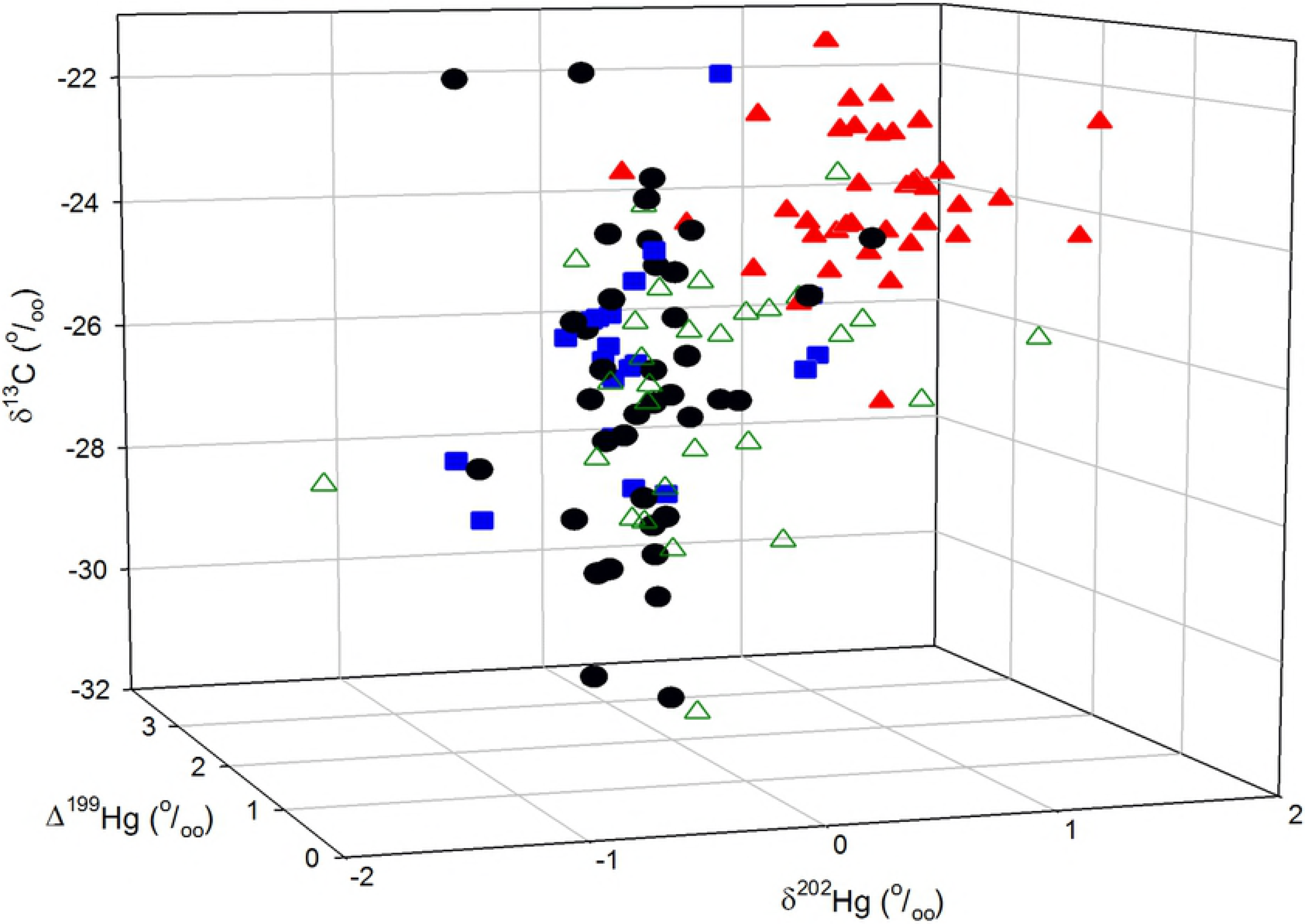
Three-dimensional plot showing mass-dependent fractionation (MDF, as δ^202^Hg) and mass independent fractionation (MIF, as Δ199) of Hg in gull and tern eggs from Egg Island and Mamawi Lake. Also shown are lipid-corrected δ^13^C values in eggs. Samples shown here were collected in 2009, 2011, 2012, or 2014. Symbols: circles are Caspian Terns, squares are Common Terns, closed triangles are California Gulls, open triangles are Ring-billed Gulls.

### Influence of riverine processes on egg THg, SSC, water clarity and Hg isotopes

Annual maximal June flow for the Athabasca River indicated large inter-year differences with flow varying approximately three-fold during the period of study (Fig. S2). The Athabasca River is unregulated by dams and annual flow is related to the volume of snowpack and glacier melt [39–41]. Mean annual THg concentrations in eggs were linearly correlated with river flow from the previous year in two of the five species/site comparisons (Egg Island: Caspian Terns n = 8, *r* = 0.90, *p* = 0.003; Common Terns n = 7, *r* = 0.89 *p* = 0.007) (Fig.4).

**Fig 4.**
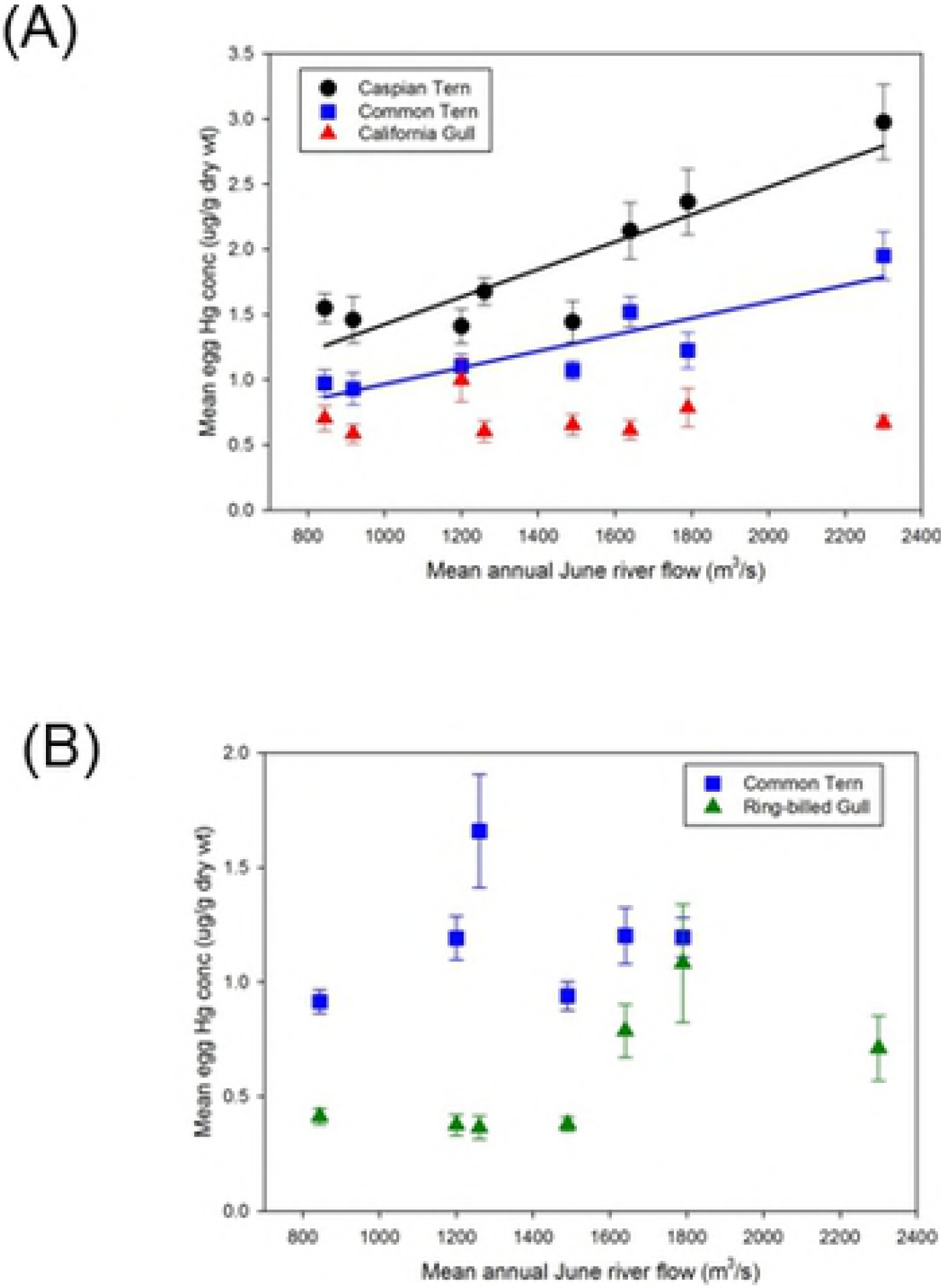
Annual mean June flow (m^3^/s) of the Athabasca River versus annual mean (± SE) THg levels (ug/g dry weight) in gull and tern eggs collected the following year. (a) Egg Island, western Lake Athabasca: Caspian Terns, Common Terns, California Gulls (b) Mamawi Lake: Common Terns, Ring-billed Gulls. Annual egg THg levels were compared to river flow in the year preceding egg collections.

However, visual examination of the data indicated that the relationship between egg THg concentrations and river flow might not be best described by a linear relationship. Instead, a threshold effect seemed more appropriate with greater egg THg levels being observed in years when river flow surpassed a mean June threshold of 1600 m^3^/s. To investigate this further, years were categorized as being low (<1600 m^3^/s) or high (≥1600 m^3^/s) flow based on June mean monthly flow. Years classified as low flow years were: 2008, 2010, 2012, 2015, 2016; high flow years were 2011, 2013, 2014. THg levels in eggs were elevated following high river flow years in the majority of species/sites examined (Fig.5). THg concentrations in eggs collected following high flow years were greater in Caspian Terns (Egg Island) (mean_low_ = 1.51 μg·g^−1^ dry wt, mean_high_ = 2.49 μg·g^−1^ dry wt; *t* = 6.85, 78 *d.f.,p* < 0.0001), Common Terns (Egg Island) (mean_low_ = 1.02 μg·g^−1^ dry wt, mean_high_ = 1.58 μg·g^−1^ dry wt; *t* = 5.45, 68 *d.f.,p* < 0.0001), and Ring-billed Gulls (Mamawi Lake) (mean_low_ = 0.38 μg·g^−1^ dry wt, mean_high_ = 0.86 μg·g^−1^ dry wt; *t* = 5.60, 81 *d.f.,p* < 0.0001). THg concentrations in eggs from these species/sites were on average 82% greater in high flow versus low flow years. Eggs from Egg Island California Gulls (*t* = 0.24, 79 *d.f., p* = 0.81) and Mamawi Lake Common Terns (*t* = 0.19, 53 *d.f., p* = 0.85) did not show a significant difference in THg levels between low and high flow years. However, exclusion of the anomalously high 2009 egg THg data for Mamawi Lake Common Terns (highest mean δ^15^N value observed that year may have indicated possible influence of dietary differences) resulted in detection of elevated THg levels in high flow years (mean_low_ = 0.98 μg·g^−1^ dry wt, mean_high_ = 1.20 μg·g^−1^ dry wt; *t* = 2.74, 43 *df, p* = 0.01).

**Fig 5.**
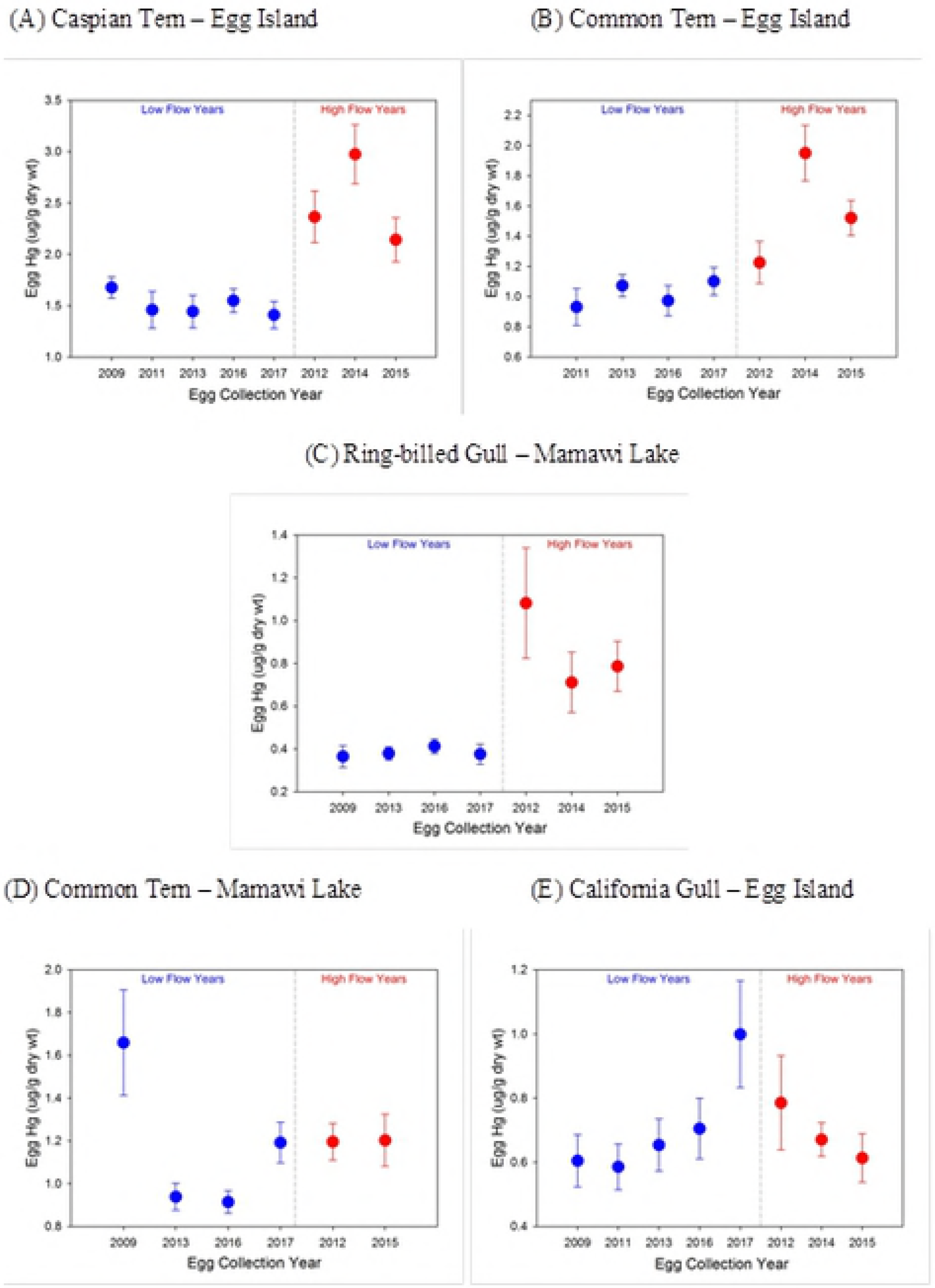
Annual mean (± SE) THg levels (ug/g dry weight) in eggs collected following years of low and high flow in the Athabasca River. Low flow years were categorized as those having a maximal annual flow of less than 1600 m^3^/s. (a) Egg Island Caspian Terns (b) Egg Island Common Terns (c) Mamawi Lake Ring-billed Gulls (d) Mamawi Lake Common Terns (e) Egg Island California Gulls.

Athabasca River flow affected the extent of sediment plumes entering Lake Athabasca (Fig.6). Furthermore, lake SSC concentrations were greater in a high flow year (2011) versus a low flow year (2010) (Fig.S3). Differences in SSC were likely responsible for inter-year differences in secchi depth in western Lake Athabasca as water clarity was lower at sites with higher SSC, particularly at sites closer to the mouth of the Athabasca River (Fig.S3). Decreased water clarity was evident in the high flow year.

**Fig 6.**
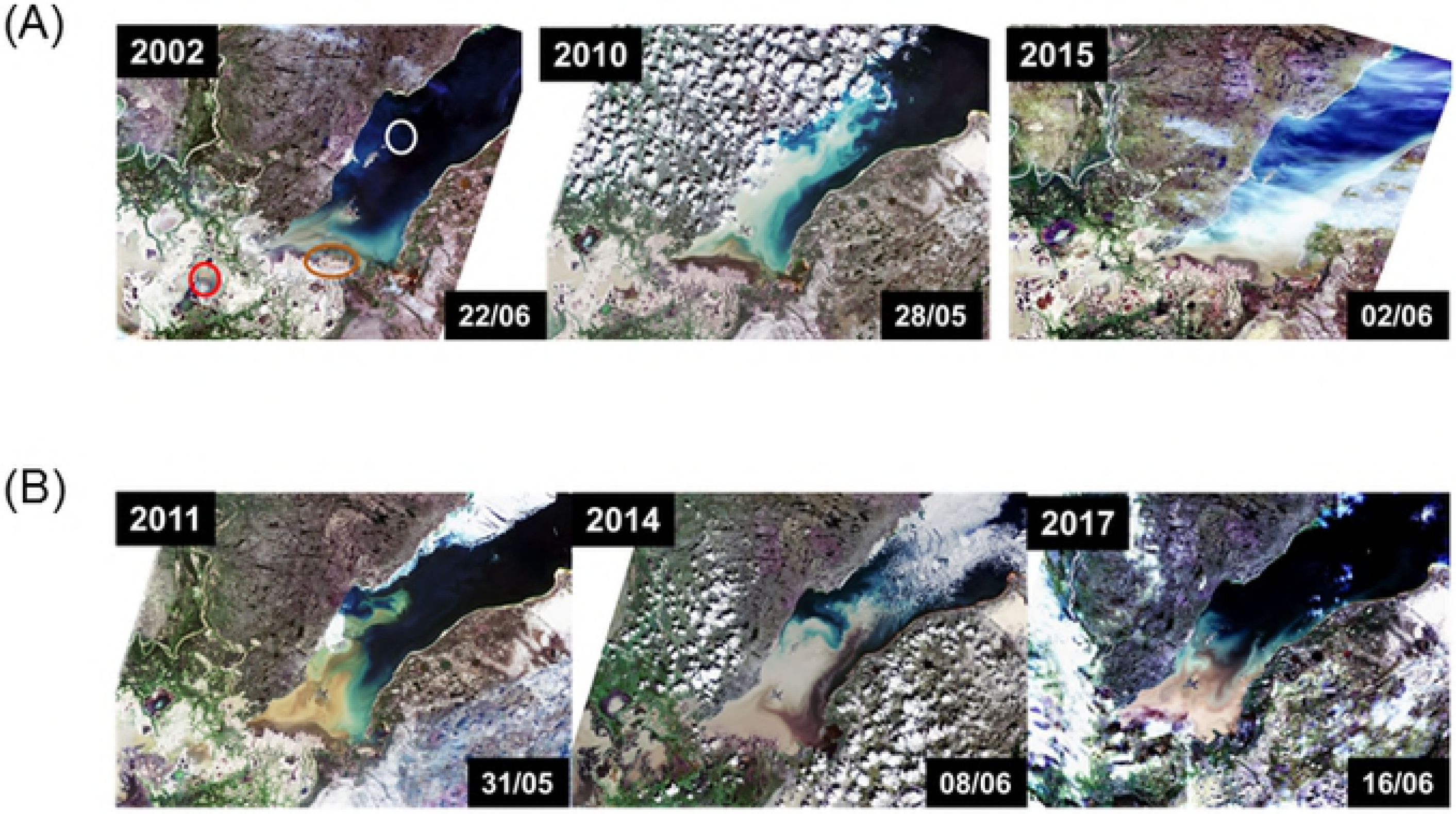
True color Landsat images of the PAD and western Lake Athabasca during years of low (<1600 m^3^/s June flow) and high (≥1600 m^3^/s June flow) Athabasca River flow. In high flow years, there is a notable increase in the extent of the brown sediment plume entering the lake. 2015 image is obscured to some extent by white cloud cover but sediment is clearly visible. Red and white circles indicate the locations of Mamawi Lake and Egg Island, respectively. Brown oval indicates the Athabasca River mouth. Dates of image acquisition are shown.

For each species, differences in egg Δ^199^Hg values were observed following years of low and high Athabasca River flow (Table 1). Statistical comparison of mean Δ^199^Hg values in eggs collected following low flow years versus high flow years indicated higher Δ^199^Hg values following low flow years in California Gulls (*t* (67) = 3.50, *p* < 0.001), Caspian Terns (*t* (38) = 2.16, *p* = 0.04), Common Terns (*t* (19) = 2.10, *p* = 0.049), Ring-billed Gulls (*t* (55) = 3.18, *p* = 0. 002). Similarly, egg Δ^201^Hg values were greater following low flow years than high flow years; California Gulls (*t* (67) = 2.62, *p* = 0.01), Caspian Terns (*t* (38) = 2.16, *p* = 0.04), Common Terns (*t* (19) = 2.68, *p* = 0.02), Ring-billed Gulls (*t* (55) = 5.04, *p* < 0.001) (data not shown). Δ^199^Hg and Δ^201^Hg values in eggs of all species were correlated (Δ^199^Hg = 0.22 + 1.07 * Δ^201^Hg, n = 187, *r* = 0.89, *p* < 0.001). Based upon inter-specific differences between gulls and terns in egg δ^13^C values (both California and Ring-billed Gulls differed from one or both tern species) and δ^202^Hg (California Gulls differed) separate regression analyses were completed for terns and gulls. For gulls, Δ^199^Hg = 0.36 + 0.89 * Δ^201^Hg, n = 126, *r* = 0.81, *p* < 0.001, and for terns, Δ^199^Hg = 0.09 + 1.25 * Δ^201^Hg, n = 61, *r* = 0.99, *p* < 0.001. δ^202^Hg values in eggs only differed between low and high flow categories for Common Terns (*t* (19) = 3.12, *p* < 0.01), other species showed no differences between low and high flow (t-tests, p > 0.3) (Table 1).

**Table 1.**
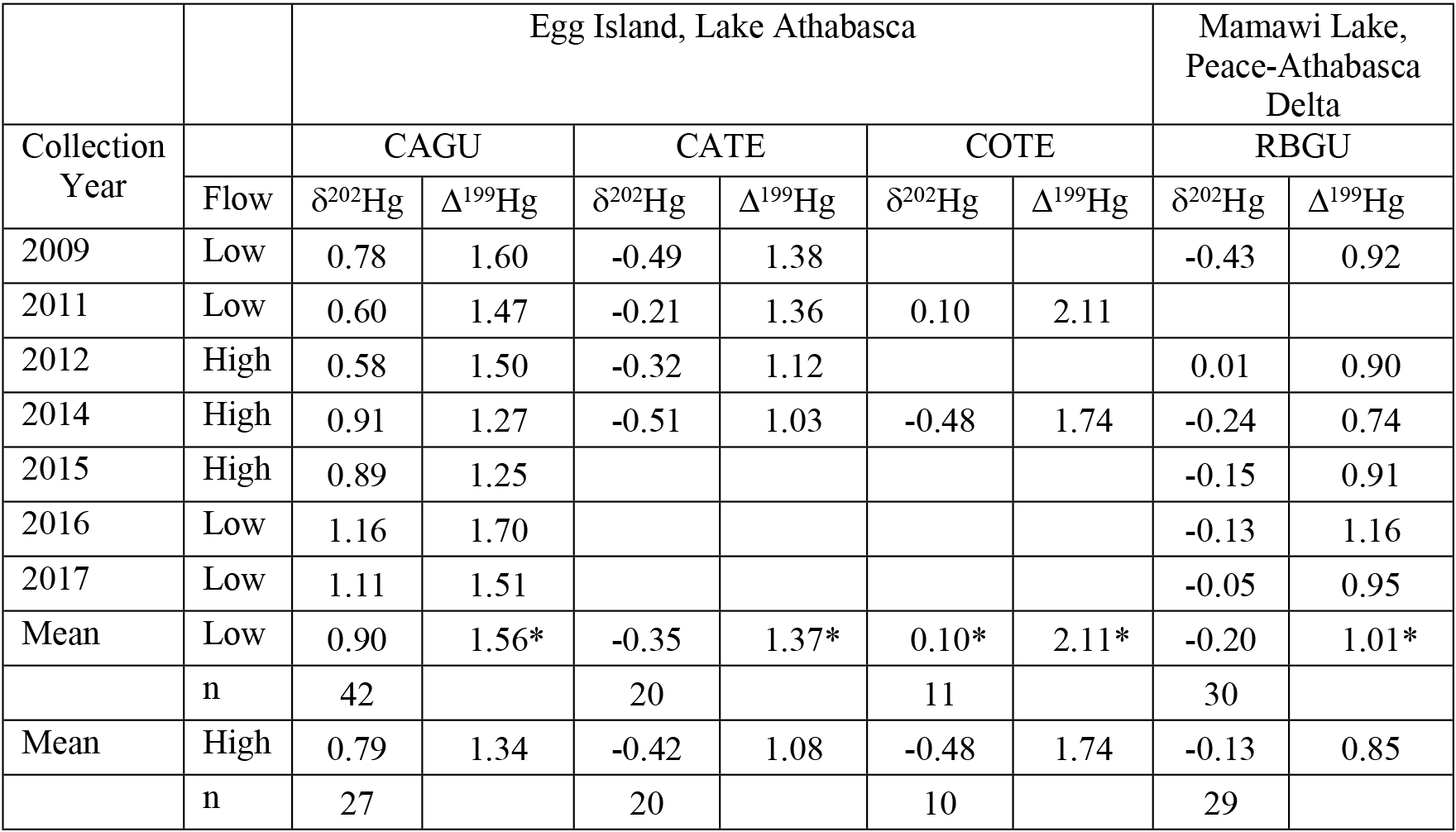
Annual mean (‰) mass dependent fractionation (MDF, δ^202^Hg) and mass-independent fractionation (MIF, Δ^199^Hg) of Hg isotopes in aquatic bird eggs. California Gulls (CAGU), Caspian Terns (CATE), Common Terns (COTE), and Ring-billed Gulls (RBGU) collected from Egg Island and Mamawi Lake. Athabasca River flow in the year preceding egg collections was categorized as low or high (≥ 1600 m^3^/s in June) for each year. MDF and MIF values were compared between flow categories for each species. * indicates statistically significant differences (t-test, *p* < 0.05) between flow categories. n is the number of egg samples included in each category.

| Collection<br>Year | Flow | Egg Island, Lake Athabasca |  |  |  |  |  | Mamawi Lake,<br>Peace-Athabasca<br>Delta |  |
| --- | --- | --- | --- | --- | --- | --- | --- | --- | --- |
|  |  | CAGU |  | CATE |  | COTE |  | RBGU |  |
| | | $\delta^{202}\text{Hg}$ | $\Delta^{199}\text{Hg}$ | $\delta^{202}\text{Hg}$ | $\Delta^{199}\text{Hg}$ | $\delta^{202}\text{Hg}$ | $\Delta^{199}\text{Hg}$ | $\delta^{202}\text{Hg}$ | $\Delta^{199}\text{Hg}$ |
| 2009 | Low | 0.78 | 1.60 | -0.49 | 1.38 |  |  | -0.43 | 0.92 |
| 2011 | Low | 0.60 | 1.47 | -0.21 | 1.36 | 0.10 | 2.11 |  |  |
| 2012 | High | 0.58 | 1.50 | -0.32 | 1.12 |  |  | 0.01 | 0.90 |
| 2014 | High | 0.91 | 1.27 | -0.51 | 1.03 | -0.48 | 1.74 | -0.24 | 0.74 |
| 2015 | High | 0.89 | 1.25 |  |  |  |  | -0.15 | 0.91 |
| 2016 | Low | 1.16 | 1.70 |  |  |  |  | -0.13 | 1.16 |
| 2017 | Low | 1.11 | 1.51 |  |  |  |  | -0.05 | 0.95 |
| Mean | Low | 0.90 | 1.56* | -0.35 | 1.37* | 0.10* | 2.11* | -0.20 | 1.01* |
|  | n | 42 |  | 20 |  | 11 |  | 30 |  |
| Mean | High | 0.79 | 1.34 | -0.42 | 1.08 | -0.48 | 1.74 | -0.13 | 0.85 |
|  | n | 27 |  | 20 |  | 10 |  | 29 |  |

Analysis of epiphytic lichen samples from the Fort McMurray region indicated relatively little spatial variation in Δ^199^Hg (mean = −0.46 ± 0.15‰, range −0.20 to −0.77‰) or Δ^201^Hg (mean = −0.54 ± 0.17‰, range −0.18 to −0.86) values. Lichen Δ^199^Hg values were used to estimate baseline MIF of Hg isotopes in atmospherically deposited Hg, an important source of Hg entering the Athabasca River. Lichen Δ^199^Hg values measured here were similar to those in abiotic environmental compartments in the oil sands region, i.e. surface soil (overburden) and road material, bitumen and mined oil sand, processed oil sand, and Athabasca River oil sand [27]. Using egg MIF data for terns from Egg Island and Mamawi Lake, the proportion of MeHg photodegraded was estimated using a modified equation from Bergquist and Blum [21].

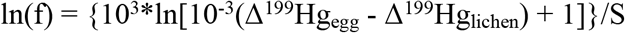

where f = fraction of MeHg photodegraded, Δ^199^Hg_egg_ = Δ^199^Hg in individual eggs, Δ^199^Hg_lichen_ = average Δ^199^Hg in lichen samples (−0.46), S = −7.82 (from the 10 mg/L dissolved organic carbon (DOC) photochemical demethylation experiment in Bergquist and Blum [21]; June 2015 DOC values in Lake Athabasca ranged from 4-7 mg/L). The amount of MeHg photodegraded was estimated to be 21.6 ± 5.5% (mean ± SD).

## Discussion

In this study, annual differences in oil sands production, bird diets, and extent of forest fires had little impact on THg levels in eggs of aquatic birds. Conversely, THg concentrations in eggs were consistently influenced by processes associated with the Athabasca River. For the majority of species/site analyses, higher egg THg concentrations were observed in eggs laid following years of high river flow. The threshold effect of river flow on egg THg levels was consistent with Long and Pavelsky’s [16] results demonstrating the influence of river flow on SSC in Lake Athabasca. Enhanced movement of sediment-associated Hg into downstream ecosystems following high river flow may be a critical factor regulating Hg availability in downstream biota. Studies in freshwater rivers have demonstrated the importance of seasonal events on the fate of Hg. In some cases, the annual export of Hg from a watershed can be determined by a single high flow event [42]. Hence, flow was likely of critical importance in moving contaminated sediments to downstream areas with resultant impacts on the bioavailability of Hg to wildlife such as birds.

Two processes may have contributed to higher egg THg levels following years of high river flow. The first stems from the possibility that in high flow years, light attenuation associated with high water SSC may have reduced the amount of photochemical degradation of MeHg. This could have resulted in an increase in the amount of MeHg available for uptake into foodwebs elevating MeHg exposure of laying females with resultant increases in egg THg concentrations. This mechanism cannot likely account for the inter-year differences in egg THg concentrations associated with river flow because only 22% of the MeHg in the tern species was estimated to have been photodegraded. The Athabasca River downstream of Fort McMurray is characterized by high SSC which reduces water transparency and possibly limits the scope for photochemical degradation of MeHg to occur. Despite the fact that photochemical degradation of MeHg was not likely responsible for inter-year fluctuations in egg THg concentrations, interyear variability in MIF of Hg isotopes was useful in linking river sources of Hg to Hg accumulated in bird eggs. Inter-year differences in egg Δ^199^Hg and Δ^201^Hg values between years of low and high flow provided evidence that riverine processes not only regulated the amount of Hg to which birds were exposed but also the isotopic composition of that Hg.

The slope (1.25) of the Δ^201^Hg/Δ^199^Hg regression for terns was similar to the value associated with photoreduction of MeHg (1.3) indicating that MeHg photoreduction was likely responsible for Δ^199^Hg and Δ^201^Hg values observed in eggs of those species. However, for gulls, a lower slope (0.89) was observed which was more similar to that associated with the photochemical reduction of Hg^2+^ (slope = 1.0, [21]). Tsui et al. [19] reported similar Hg isotope patterns in biota associated with aquatic (slope = 1.20 in benthic invertebrates, slope = 1.24 in trout) versus terrestrial food webs (slope = 1.05). They hypothesized that the mechanisms underlying MIF of Hg differed between aquatic and terrestrial ecosystems. In terrestrial systems, Hg^2+^ may be extensively photoreduced before the non-photoreduced Hg^2+^ undergoes methylation. Hence, differences in Hg sources, pathways of exposure, and environmental Hg processing, could explain the differences in egg Hg trends detected in California Gulls (i.e. detection of a fire signal) versus the other species. Increased use of terrestrial resources by gulls was supported by both the egg Hg MDF (δ^202^Hg) and δ^13^C data. Tsui et al. [19] reported higher δ^202^Hg values in organisms associated with terrestrial systems; similar to what was observed for California Gulls in this study. Carbon isotope results corroborated this interpretation as egg δ^13^C values in terns were more negative than those measured in gulls. This was consistent with what would be expected based upon differences in δ^13^C values of food originating from freshwater and terrestrial ecosystems [43] and suggested that gulls were incorporating terrestrial foods into their diets in addition to aquatic prey. These results reflected the well-characterized, more omnivorous diet of gulls (see [44] for an example). However, all of the bird species, including gulls, exhibited MIF of Hg that reflected the importance of aquatic processes in regulating egg Hg isotope values. Consistent differences in Δ^199^Hg values between low and high flow year categories for all bird species indicated that all of them were linked to aquatic ecosystem processes. It is just possible that gulls may also show connections with terrestrial processes as well.

Here, we do not provide direct evidence linking Hg levels in eggs to oil sands sources. However, previous studies have demonstrated that oil sands developments are a source of Hg to the local environment [3, 4]. For example, Hg in snow was 5.6 times greater within 50 km of oil sands processing facilities than outside that area [3]. Hg levels in water sampled in summer were three times greater downstream of areas disturbed by oil sands development than at upstream locations situated in the mineable oil sands region [3]. Water Hg concentrations further downstream in the Athabasca River and PAD were two times greater than these upstream concentrations [3]. Elevated levels of Hg in abiotic environmental matrices are also reflected in biota. For example, THg concentrations in prey fish sampled from the Athabasca River in the surface mineable oil sands region were five times greater than fish collected from the Athabasca River upstream of the oil sands [15]. THg levels in prey fish collected in the PAD and Lake Athabasca were two to four times greater than fish from the upstream Athabasca River site [15]. THg levels in eggs of California Gulls and Herring Gulls (*Larus argentatus*) were two and three times greater, respectively, at Egg Island (Lake Athabasca) than at Namur Lake, an inland lake isolated from the Athabasca River but in close proximity (~60 km west) to open-pit oil sands mines [15]. Taken together, these results indicate that the oil sands are a source of Hg to the environment and that biota inhabiting waters in or downstream of the oil sands have higher Hg levels. To understand this further, an integrated research program involving Hg measurements in air, water, land, and biota is required to assess the relative importance of oil sands sources as a contributor to the overall Hg budget of the Athabasca River and downstream ecosystems. With respect to bird eggs, we can begin to predict expected Hg levels based upon river flow. For example, in 2017, mean June river flow was high (~1590 m^3^/s), hence egg THg levels in 2018 are also predicted to be high. Eggs are currently being analyzed to test this hypothesis.

Until now, uncertainty surrounded the degree to which Hg may be transported by the Athabasca River to ecosystems far downstream. Following spring melt of the snowpack in the Athabasca River watershed, it is likely that some of the snow-associated mercury finds its way into the river. This is particularly true in the spring when frozen soils may limit infiltration of runoff into soil leading to efficient Hg export to the Athabasca River from overland sources [45, 46]. Furthermore, during high flow years, Athabasca River sediments are mobilized and transported to downstream environments such as the PAD and western Lake Athabasca [16]. River systems can convey 90% of their total heavy metal load via sediment transport [17, 42,
46]. Hence, it is highly likely that Athabasca River sediments are transporting Hg to areas far
downstream. This hypothesis was supported by Long and Pavelsky’s [37] data and by satellite images showing inter-year differences in sediment plumes into Lake Athabasca. Aquatic bird egg THg concentrations and Hg isotope data indicate that this Hg is being incorporated into downstream food webs.

The importance of the current study lies in its highlighting the degree to which local inputs of Hg to the river via atmospheric releases, mobilization associated with land disturbance, or dust/leakage from tailings ponds (but see [47]) will not remain confined to the local receiving environment. Hg from these sources will be transported long distances via the river to sensitive downstream ecosystems, e.g. PAD, WBNP, that are recognized for their unique, world-class ecological characteristics. New surface-mine oil sands projects are being proposed that will bring development much closer (~30 km) to the southern boundary of the PAD/WBNP. This will likely increase Hg inputs into the local environment through Hg mobilization stemming from further land development and atmospheric Hg releases. The zone of atmospheric deposition may encompass southern parts of the PAD/WBNP based upon the size of depositional zones previously characterized for oil sands operations [3, 4]. Hence, further oil sands development may result in increased delivery of Hg to downstream/downwind areas. The impacts of multiple stressors (including oil sands) on WBNP have resulted in it being investigated as an UNESCO World Heritage Site in Danger. Cumulative impact assessment of existing and proposed oil sands mining projects needs to consider potential Hg impacts on wildlife and humans inhabiting this globally-recognized area of ecological importance.

## SUPPORTING INFORMATION

**Figure S1** shows temporal trends in surface-mineable oil sands production. **Figure S2** shows annual trends in Athabasca River June flow. **Figure S3** contrasts suspended sediment concentrations and water transparency in Lake Athabasca in years of low versus high river flow. **Table S1** and **Table S2** tabulate stable carbon and nitrogen isotope data by species, site and year, respectively.

## Acknowledgments

The Mikisew Cree First Nation (MCFN), the Athabasca Chipewyan First Nation, and Métis Local 125 are thanked for their support and for granting permission for collections on their traditional lands. J. Marten, L. McKay (MCFN); A. Caron, S. Dolgova, L. Shutt, D.V.C. Weseloh, M. Zanuttig (ECCC); D. Campbell, Q. Gray, R. Kindopp, J. Lankshear, L. Patterson, J. Straka (Parks Canada Agency); B. Maclean, G. Paterson and E. Seed assisted with egg collections. Fort Chipewyan Community-based Monitoring Program staff provided prey fish. J. Chapman (Carleton University) sorted prey fish according to species. J. Chételat, L. Mundy, C. Boutin, D. Carpenter, H. Gill, and P. Thomas (ECCC) supplied lichen samples for Hg isotope analysis. Tissue Processing Laboratory staff, National Wildlife Research Centre, are thanked for their expert preparation of samples. S. Dolgova, E. Porter (ECCC) conducted the egg mercury analyses. W. Abdi, G.G. Hatch Stable Isotope Laboratory, University of Ottawa, is thanked for stable nitrogen and carbon isotope analysis. Stable mercury isotope analysis was conducted by B. Georg, Water Quality Centre, Trent University; P. Dillon facilitated these analyses. V. Wynja (ECCC) sourced the Landsat images and the United States Geological Survey is thanked for making the images available. This research was funded by the Joint Canada-Alberta Oil Sands Monitoring Program.

## Supplementary Information

### Captions

S1_Table. Annual mean (± 1 SD) δ^15^N values (‰) in eggs of California Gulls (CAGU), Caspian Terns (CATE), Common Terns (COTE), and Ring-billed Gulls (RBGU) collected from Egg Island and Mamawi Lake. At each site, inter-year differences in species-specific δ^15^N values were evaluated using ANOVA or Kruskal-Wallis/Dunn’s tests. Superscript letters indicate statistically significant differences (*p* < 0.05) between years. Means with the same letter are not different. n is the number of samples analyzed for each species at each site.

S2_Table. Annual mean (± 1 SD) δ^13^C values (‰) in eggs of California Gulls (CAGU), Caspian Terns (CATE), Common Terns (COTE), and Ring-billed Gulls (RBGU) collected from Egg Island and Mamawi Lake. δ^13^C values were adjusted for lipid content. At each site, inter-year differences in species-specific δ^13^C values were evaluated using ANOVA/Tukey’s HSD or Kruskal-Wallis/Dunn’s tests. Superscript letters indicate statistically significant differences (*p* < 0.05) between years. Means with the same letter are not different. n is the number of samples analyzed for each species at each site.

S1_Fig. Surface-mineable oil sands production (synthetic oil and bitumen) in millions of barrels per day, 1967-2016. Data are from CAPP [2]. The dashed line indicates the beginning of the period during which egg samples were collected to assess temporal trends in mercury levels.

S2_Fig. Mean June flow (m^3^/s) of the Athabasca River north of Fort McMurray (station 07DA001) from 1958-2016. The dashed line indicates the beginning of the period during which the influence of river flow on egg THg levels was investigated. Data are from Canadian Hydrographic Service (https://wateroffice.ec.gc.ca/mainmenu/historical_data_index_e.html).

S3_Fig. Suspended sediment concentrations (SSC) (mg/L) and secchi disc depth (cm) along a transect from a point (58.6724 N, −110.8477 W) at the mouth of the Athabasca River. Data for a low flow (2010) and a high flow year (2011) are shown. Data are from Long and Pavelsky [37].

